# Drought Affects the Antioxidant System and Stomatal Aperture in *Zanthoxylum bungeanum* Maxim

**DOI:** 10.1101/578294

**Authors:** Xitong Fei, Haichao Hu, Jingmiao Li, Yulin Liu, Anzhi Wei

## Abstract

When under drought, plants activate a range of self-protective responses. Among these are activation of the antioxidant system and changes in stomatal aperture. The antioxidant system can remove the reactive oxygen species produced under drought conditions and so mitigate oxidative damage. Water becomes a severely limiting resource for plants suffering drought stress, so they generally close their stomata to reduce water loss. We examined changes in the activities of the antioxidant enzymes and altered gene expression patterns in *Zanthoxylum bungeanum* plants exposed to drought by irrigation with 20% PEG6000. We also recorded changes in stomatal aperture as the drought persisted. Relationships between the antioxidant system and stomatal aperture were analyzed in relation to gene expression. The results indicate that under drought stress, POD, CAT, APX, proline, MDA and related genes all show positive responses to drought, while SOD and its genes showed a negative response. The relationship between drought duration and stomatal aperture was considered. Stomatal aperture declines exponentially as drought duration increases.

## Introduction

*Zanthoxylum bungeanum* Maxim. (common name Chinese prickly ash, family *Rutaceae*) is widely distributed in Asia [1] where it is an important economic crop. Evolution and natural selection have led the epidermis of *Z. bungeanum* to bear prickles. This species also has strong drought adaptability. The skin of *Z. bungeanum* is the source of one of the eight traditional Chinese condiments, so this plant plays a very important role in Chinese food culture. Because of its unique numbing taste, *Z. bungeanum* is difficult to replace with other seasonings [2]. It has become an important component of the diet in various parts of Asia, especially in China. *Zanthoxylum bungeanum* and pepper become best companions and are together an important part of the ‘hot pot’ culture. In addition to its use as a food seasoning, the skin of *Z. bungeanum* also contains chemical components showing proven medicinal properties, including bactericidal [3], insecticidal [4], antioxidant [2] and topical anesthetic [5].

Drought stress can cause a series of physiological and molecular reactions in plants, which seriously affect normal growth. Thus, drought can cause imbalances in cellular reactive oxygen species (ROS), it can also upset cell membrane lipid peroxidation and it can damage cell and organelle membranes. Excessive ROS have toxic effects on plants. Irrigated agriculture is not yet general, so drought remains one of the most important unfavorable factors affecting both the yield and quality of most commercial crops. Plants have many protective responses to maintain metabolic stability and so continue life under environmental stress. Their antioxidant systems are able to produce a variety of antioxidant enzymes - including superoxide dismutase, peroxidase, antioxidant enzymes, ascorbate peroxidase - to combat the ROS produced under drought stress [6, 7]. For example, catalase can decompose H_2_O_2_ produced in plants to form water and oxygen, reducing or eliminating damage by this ROS [8]. The antioxidant system also maintains organelle stability, preventing damage to the chloroplast membrane and so stabilizing the PSII system [9]. In addition, stomata are important gas exchange organs of plants, playing irreplaceable roles in the regulation of photosynthesis, respiration, transpiration and temperature [10, 11]. The stomata are also the water-regulating organs of plants. Under drought, water conservation becomes the decisive factor for plant survival. The response of stomata to drought is also a way for plants to protect themselves. Under drought stress, plants reduce water loss by stomatal closure and so increase their ability to resist drought [12, 13].

The impact of drought on the yield and quality of *Z. bungeanum* is huge and seriously hinders the development of this industry. The detailed responses of the antioxidant system and of the stomata of *Z. bungeanum* to drought stress remain unclear. Hence a study of their behavior under drought stress, will have significance for better understating this species’ drought-resistance mechanisms. It can also provide a basis for drought-resistant breeding of *Z. bungeanum* and of related species.

## Materials and methods

### Materials

The *Z. bungeanum* seeds were collected from the Fengxian Chinese prickly ash Experimental Station of Northwest Agriculture and Forestry University. They were germinated and cultured in an artificial climate chamber at 25±2°C. Air humidity was set to 80% and the photoperiod to 16:8 h (light:dark). Three-month-old seedlings were used as experimental material. *Zanthoxylum bungeanum* seedlings were then cultured in half-strength Murashige and Skoog (MS) liquid medium containing 20% PEG6000. Leaf samples were collected and stored in liquid nitrogen after periods of 0, 3, 6, 12, 24, 36 and 48 h.

### Methods

#### Physiological index determination

The leaf samples after different periods of drought stress were used to determine antioxidant enzyme activity and malondialdehyde (MDA) and proline contents. Superoxide dismutase (SOD) activity was determined by the nitroblue tetrazolium method [7]. Peroxidase (POD) activity was determined by the guaiacol method and catalase (CAT) activity was determined by the hydrogen peroxide ultraviolet method [8]. For APX (L-ascorbate peroxidase) activity we used the method of Panchuk et al [14]. The MDA content was determined by the thiobarbituric acid (TBA) method [15]. Proline was determined by ninhydrin colorimetry [16].

#### Total RNA extraction

Total RNA of the *Z. bungeanum* samples was extracted using the TaKaRa MiniBEST Plant RNA Extraction Kit (TaKaRa, Beijing, China) following the manufacturer’s instructions. The purity and concentration of the RNA obtained were measured using NanoDrop 20000 (Thermo Scientific, Pittsburgh, PA, USA). Only samples where the OD260/280 value was 1.8-2.0 and the OD260/230 value was higher than 2.0 were used for cDNA synthesis.

#### Quantitative Real-time PCR

Primer 7.0 software (Premier, Palo Alto, CA, USA) was used to design RT-qPCT primers (see Table 1). The qRT-PCR assays were carried out on a CFX96 Real-Time PCR Detection System (Bio-Rad, Hercules, CA, USA). The reaction system was of 10 μl, containing 5 μl of 2× SYBR Premix Ex Taq II (TaKaRa), 1 μl of cDNA, 1 μl of each of the upstream and downstream primers and 2 μl of ddH_2_O. *ZbUBQ* and *Zbα-EF* were used for the reference genes to correct the RT-qPCR data [17].

**Table 1.**
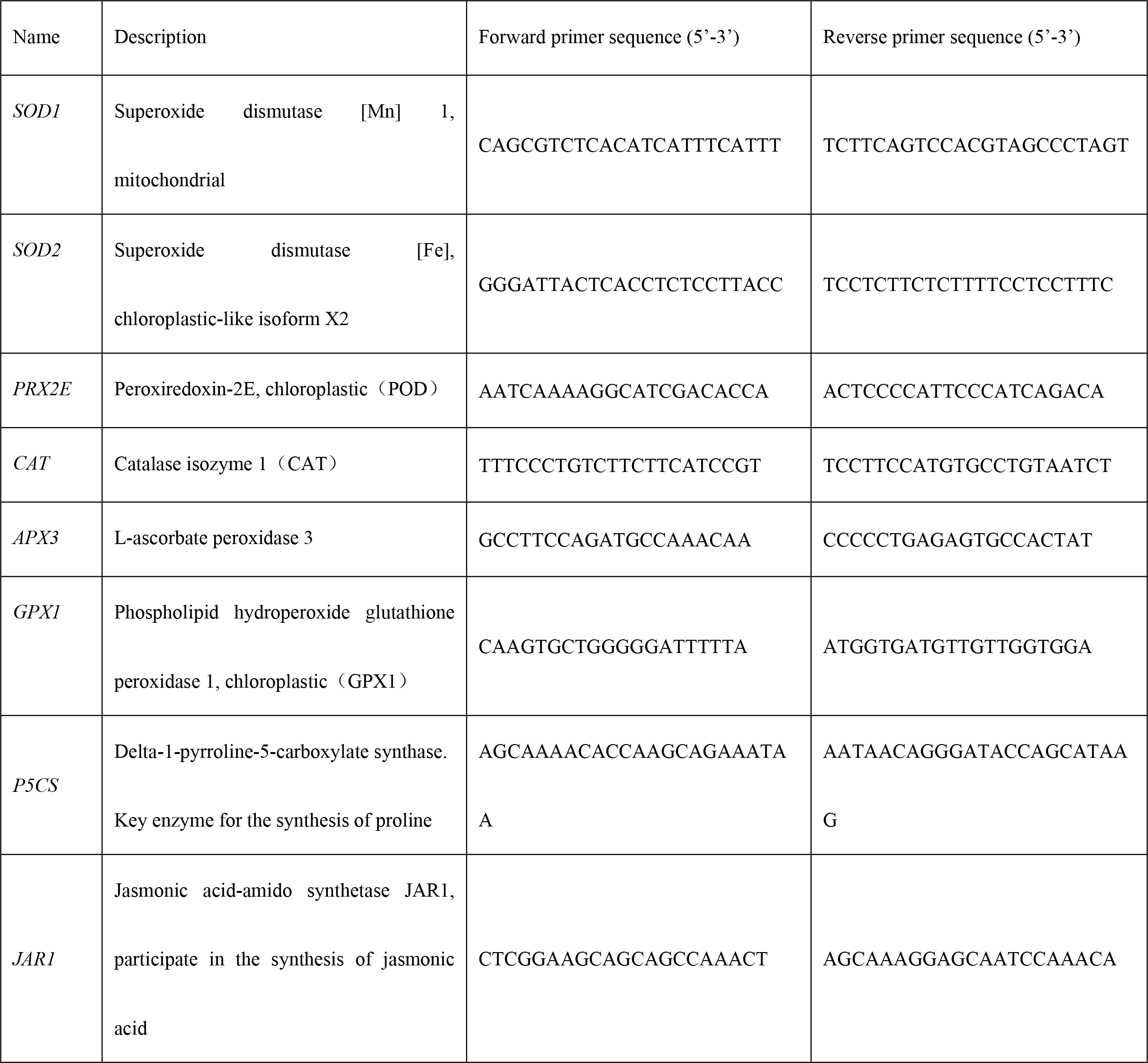
RT-qPCR primers.

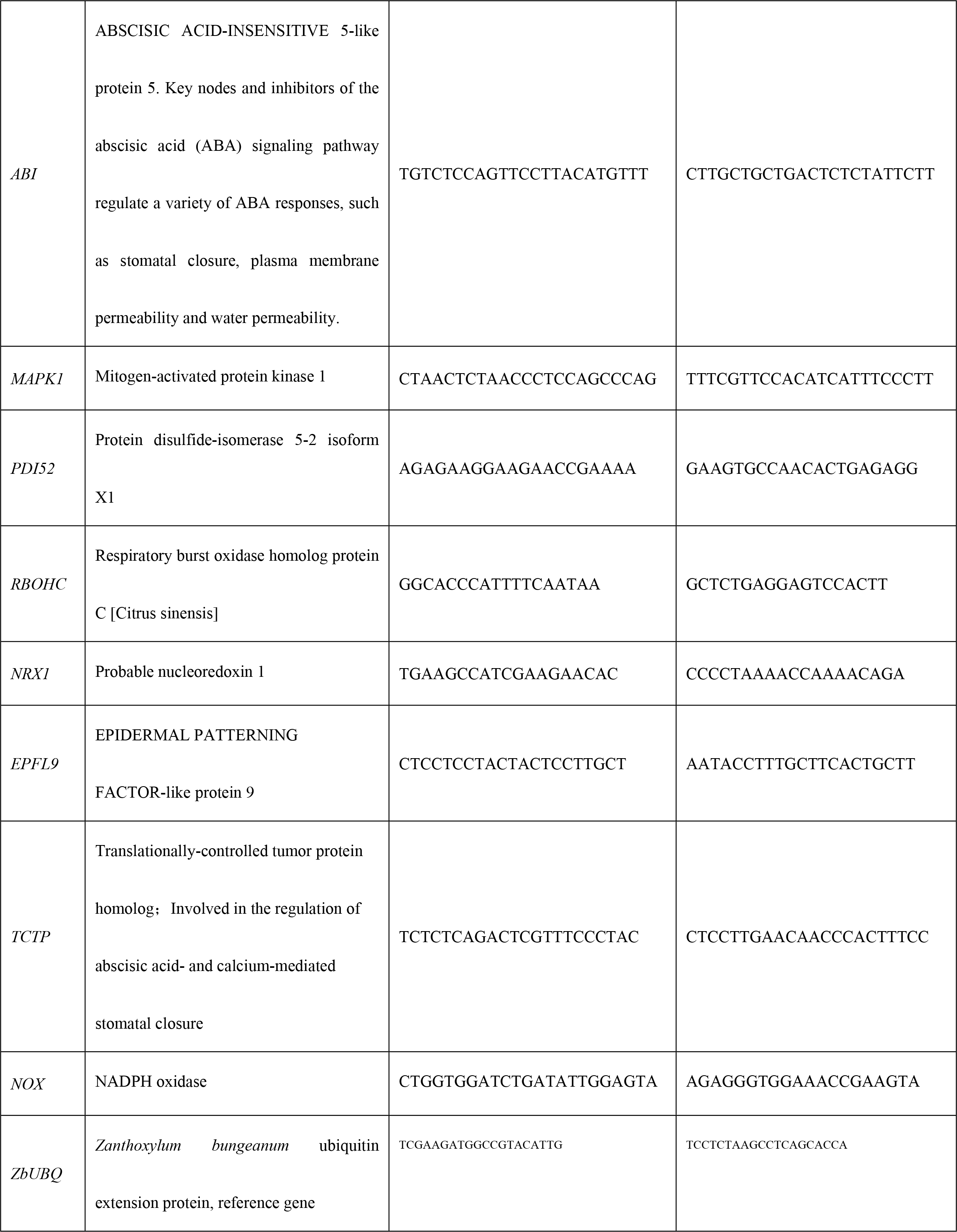

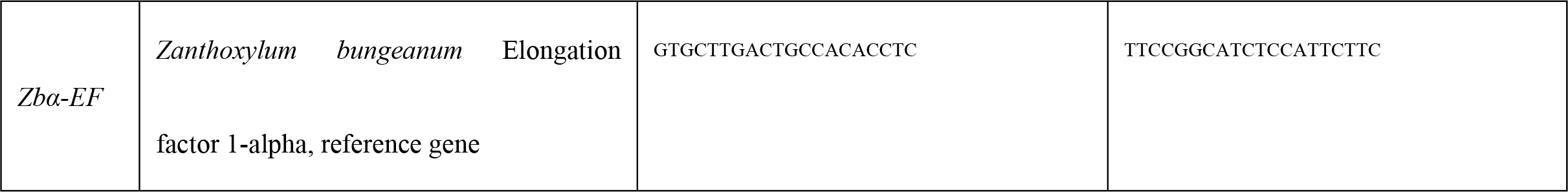

#### Stomatal observation

The nail-polish method was used to observe the state of the stomata on the leaves. Nail polish was applied to the abaxial surfaces of *Z. bungeanum* leaves after different periods of drought stress, air-dried, peeled off and examined under a light microscope (Olympus Provis AX70 microscope, Olympus Medical Systems Co., Tokyo, Japan) [18]. Thirty fields were randomly selected for observation. The numbers of stomata per unit area were calculated as stomatal density. Thirty stomata were randomly selected for each period to measure stomatal aperture.

## Results

### Physical substance detection

Changes in the levels of protective substances and genes in antioxidant system were monitored after different periods of drought stress. The results show the activities of POD, CAT and APX follow the same general pattern - their activities increased under continuing drought stress and remained at more constant high levels in the later stages (Figure 1). However, SOD activity gradually decreased with increasing duration of drought stress. The contents of proline and MDA also increased with drought duration.

**Figure 1.**
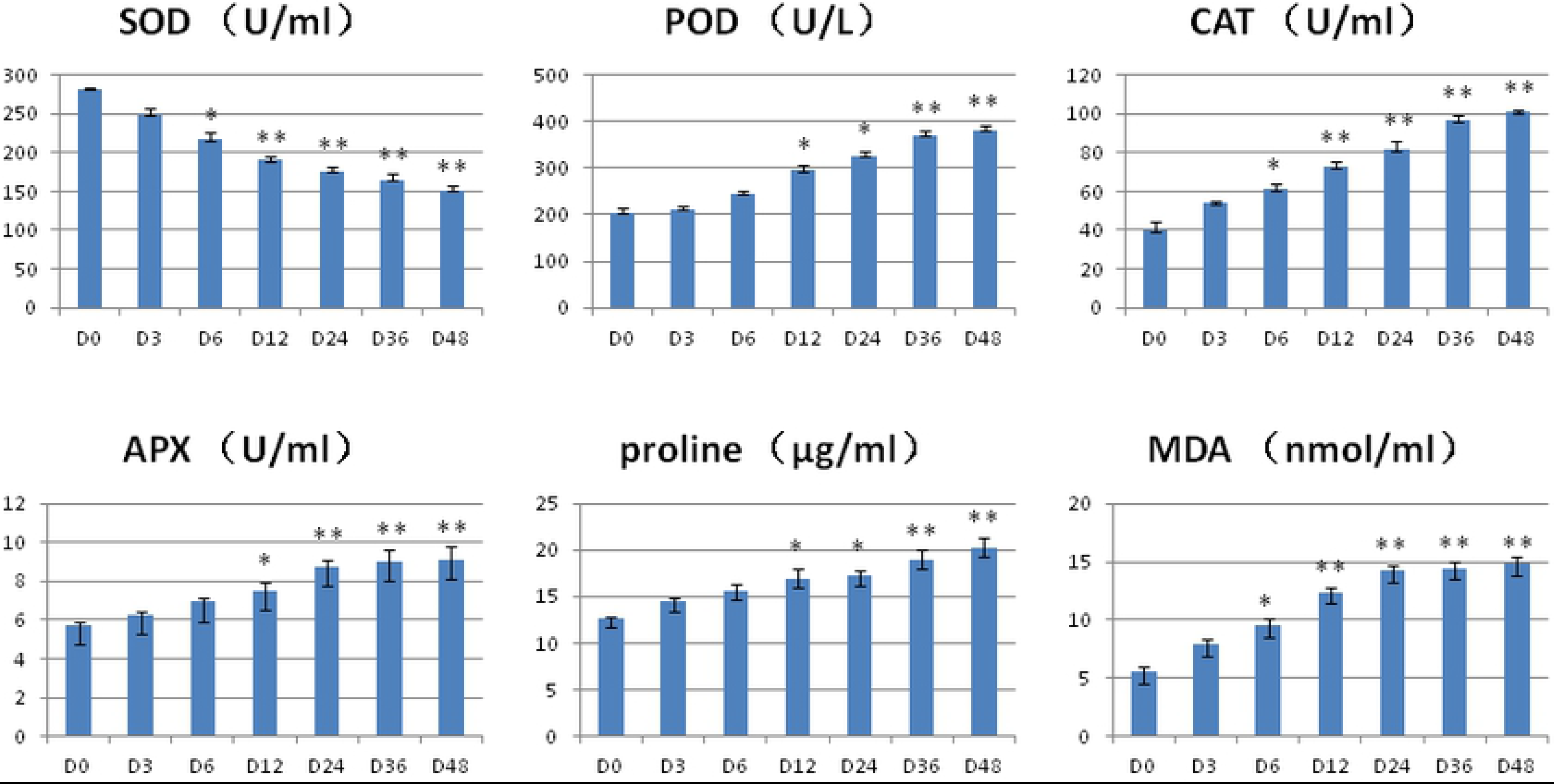
Antioxidant system enzyme activity and proline and MDA contents.

### Molecular level detection

The expression patterns of genes related to the antioxidant system were essentially consistent with the changes in the related substances. Genes such as *PRX2E*, *CAT*, *APX3*, *P5CS* and *GPX1* showed a significant increase under drought stress (Figure 2). The expression levels of the SOD gene in chloroplasts and mitochondria were monitored and we found the *SOD2* gene in the chloroplast was positively correlated with superoxide dismutase activity. The *SOD1* gene in mitochondria was negatively correlated with the activity of superoxide dismutase. It is concluded that the superoxide dismutase produced by *Z. bungeanum* under drought stress comes mainly from the chloroplast.

**Figure 2.**
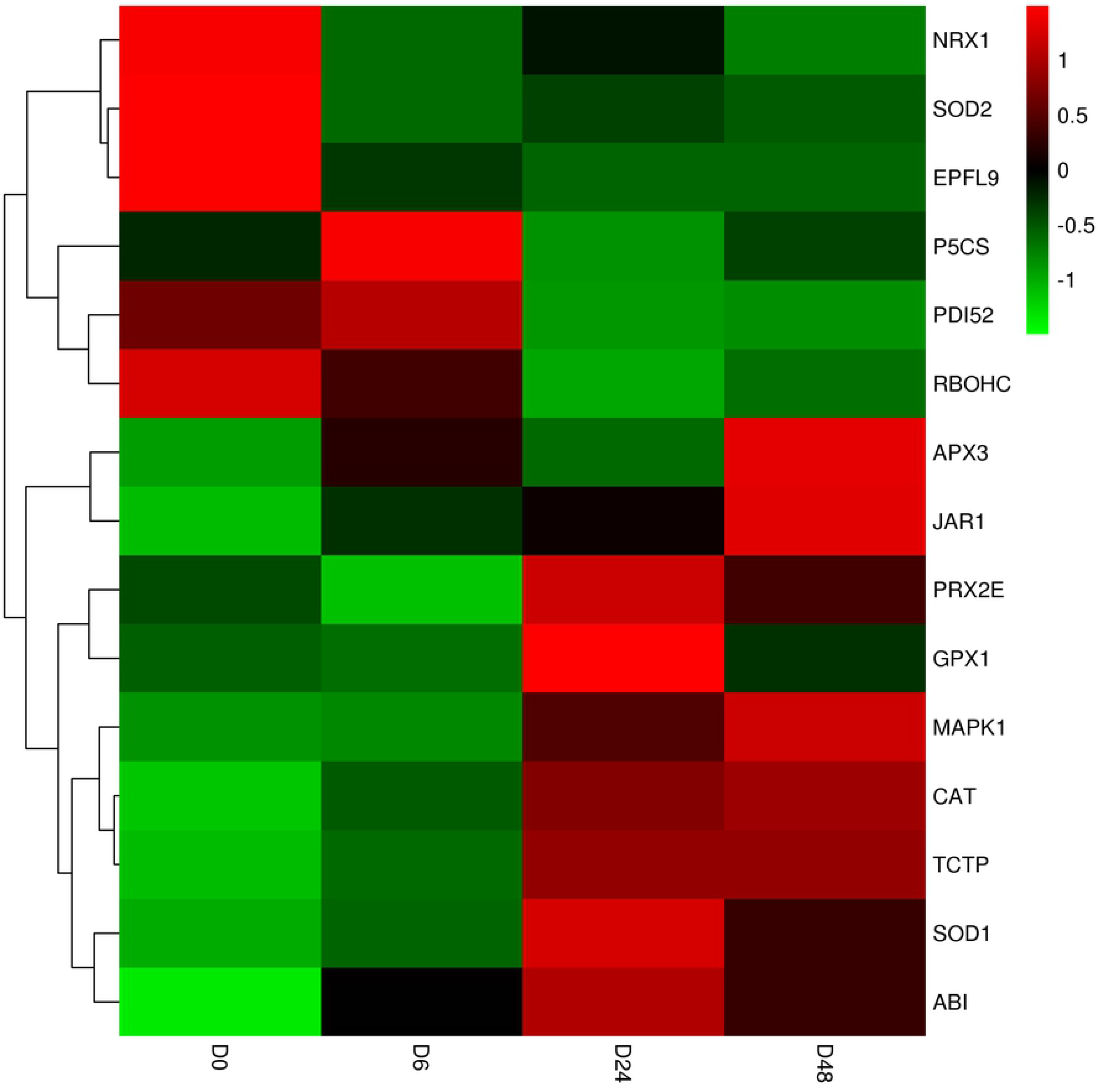
Heat map of gene expressions relating to the antioxidant system and stomatal aperture.

Some further gene expression patterns with drought stress and stomatal movement were also monitored. *P5CS* is a key enzyme in the proline synthesis process. *ABI* is a key inhibitor of the ABA signaling pathway and participates in the closure of stomata and *TCTP* is involved in ABA and calcium ion-mediated stomatal closure. In addition, the relative expression levels of *MAPK1* and *PDI52* were also up-regulated (Figure 3). At the same time, the relative expression of JAR1, a gene related to jasmonic acid synthesis, was also up-regulated under the induction of drought stress. However, other genes were inhibited, such as *RBOHC*, *NRX1*, *EPFL9*.

**Figure 3.**
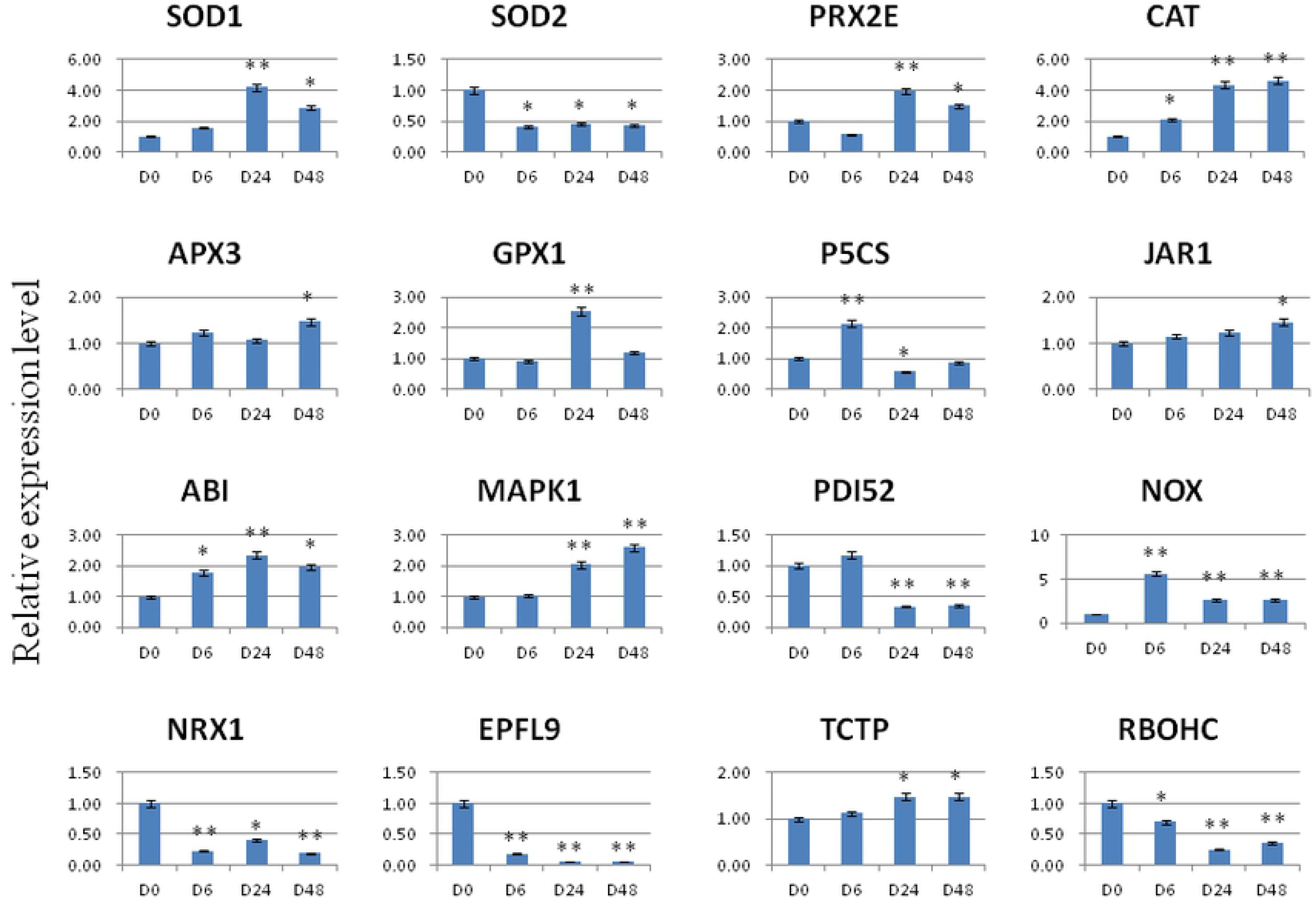
Relative expression levels of genes related to antioxidant system and stomatal aperture under continuing drought stress.

### Stomatal opening

The numbers of stomata on the abaxial leaf surfaces of *Z. bungeanum* were counted and stomatal dimensions were measured. Average stomatal density was found to be 850.87 mm^−2^. Average guard cell length was 13.51 μm and width was 8.95 μm. Average length of the stomatal pores was 17.03 μm and their maximum width (aperture) was 7.45 μm (see Figure 4).

**Figure 4.**
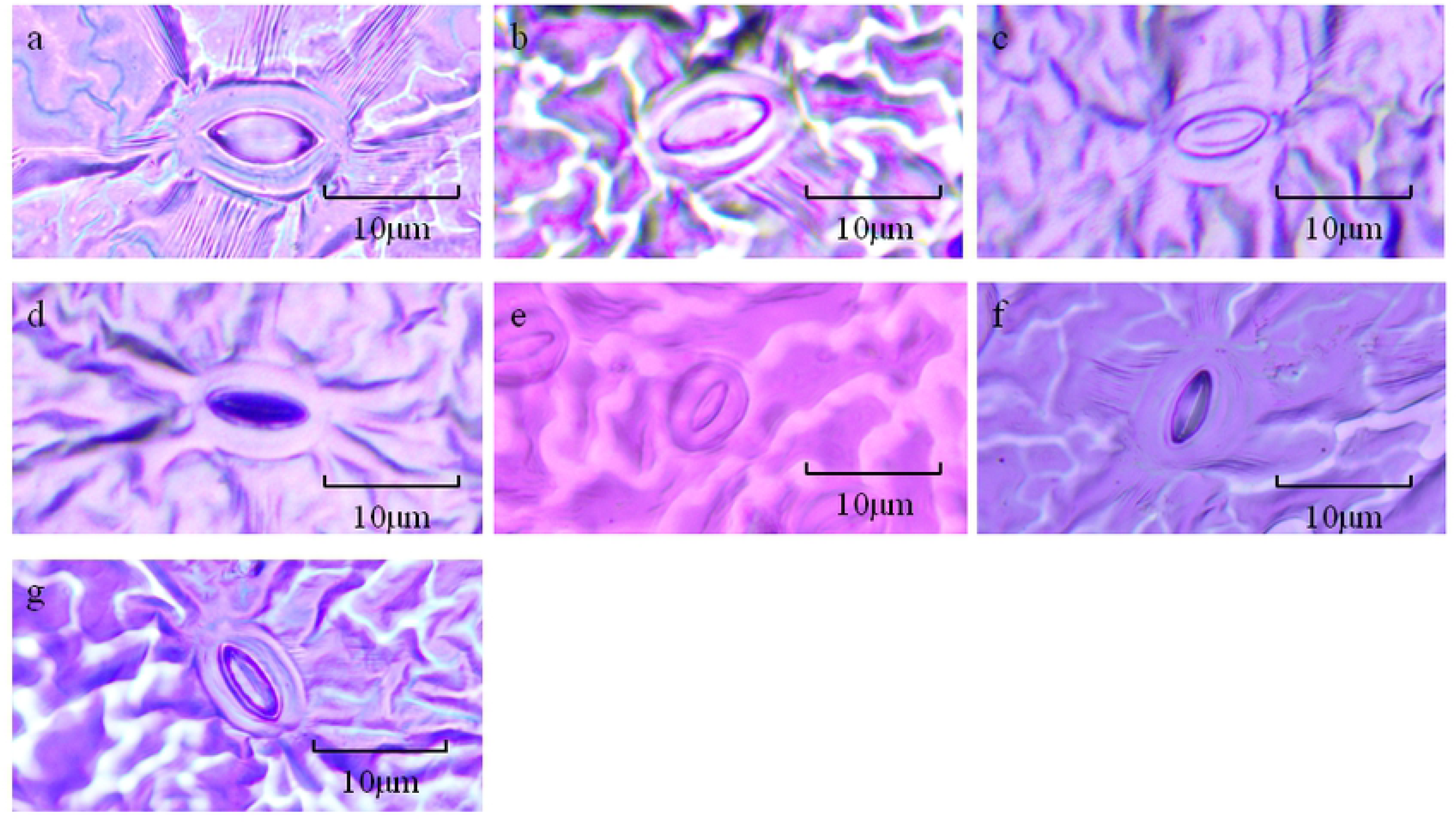
Stomatal movement of *Zanthoxylum bungeanum* under drought stress.

The morphology of the stomata after different durations of drought stress was recorded. As can be seen from Figure 4, the stomata responded significantly to drought stress duration. Stomatal width at different stages of drought stress was measured.

**Figure 5.**
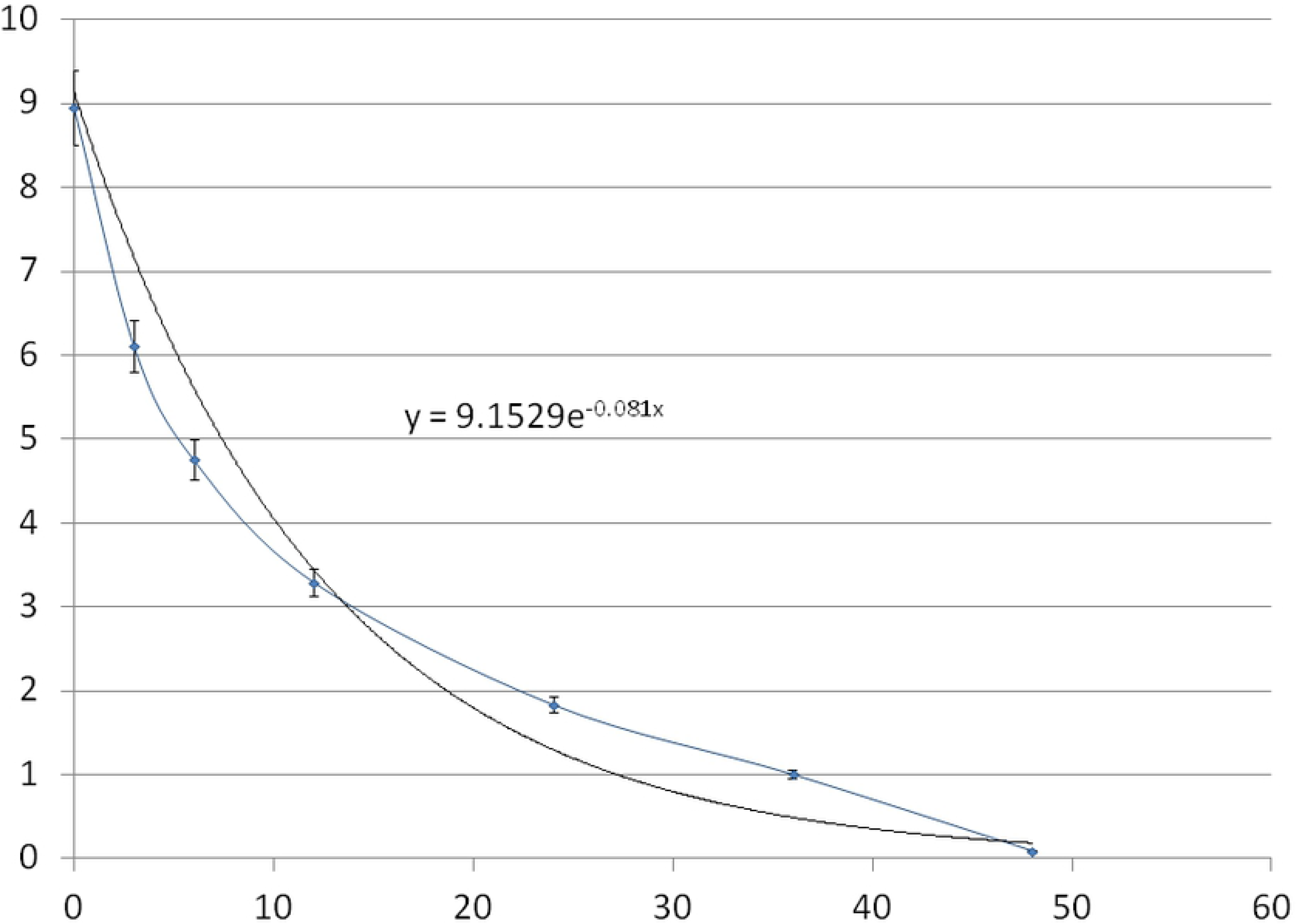
Trend of stomatal width with drought stress duration.

The results for stomatal aperture vs. drought stress duration (Figure 5) were best fitted by the exponential equation (Y=9.1529e^−0.081X^, R^2^=0.92) where the final value for slope represents the time at which the stomata are closed. With increasing drought stress duration, stomatal aperture gradually decreased and reached almost complete closure at 48 h. However, during this period, the rate of stomatal closure was decreasing. That is, the early rate of stomatal closure under drought stress was rapid, and this rate decreased with time.

## Discussion

The results show that most of antioxidant enzymes trend upward under continuing drought stress, while SOD shows a downward trend. The trend of the *SOD2* gene and SOD in the chloroplasts are consistent, which indicates the chloroplast antioxidant system was severely damaged during the drought. In addition, it can be explained that in the antioxidant system, several other antioxidant enzymes (such as POD, CAT, APX) exert major antioxidant effects. Proline and MDA increased gradually during the drought stress and may also be involved in signal transduction and protection. Studies have shown that *Barbula fallax* and pepper have similar patterns of antioxidant enzyme activity, where SOD showed a downward trend in the early stage of drought stress, while POD and CAT activities were positive in response to drought stress and increased in the early stage of drought stress [19]. In many species, the activities of antioxidant enzymes such as SOD, POD, CAT, and APX generally rise under drought. Chickpea accumulates proline and increases the activity of SOD, APX, GPX and CAT under drought stress [20]. The activities of SOD, POD and CAT in alfalfa increase significantly under drought stress [21]. This also occurs in pea [22], rice [23], Kentucky bluegrass [24] and sesame [25].

### Relationship between antioxidant signaling pathway and stomatal aperture

There are many reports on changes in antioxidant enzyme activity and stomatal morphology under drought stress but the regulation of stomata by the antioxidant system is rarely reported under drought stress. Based on our experimental results, we analyzed the regulatory relationship between the oxidative system and stomatal aperture.

**Figure 6.**
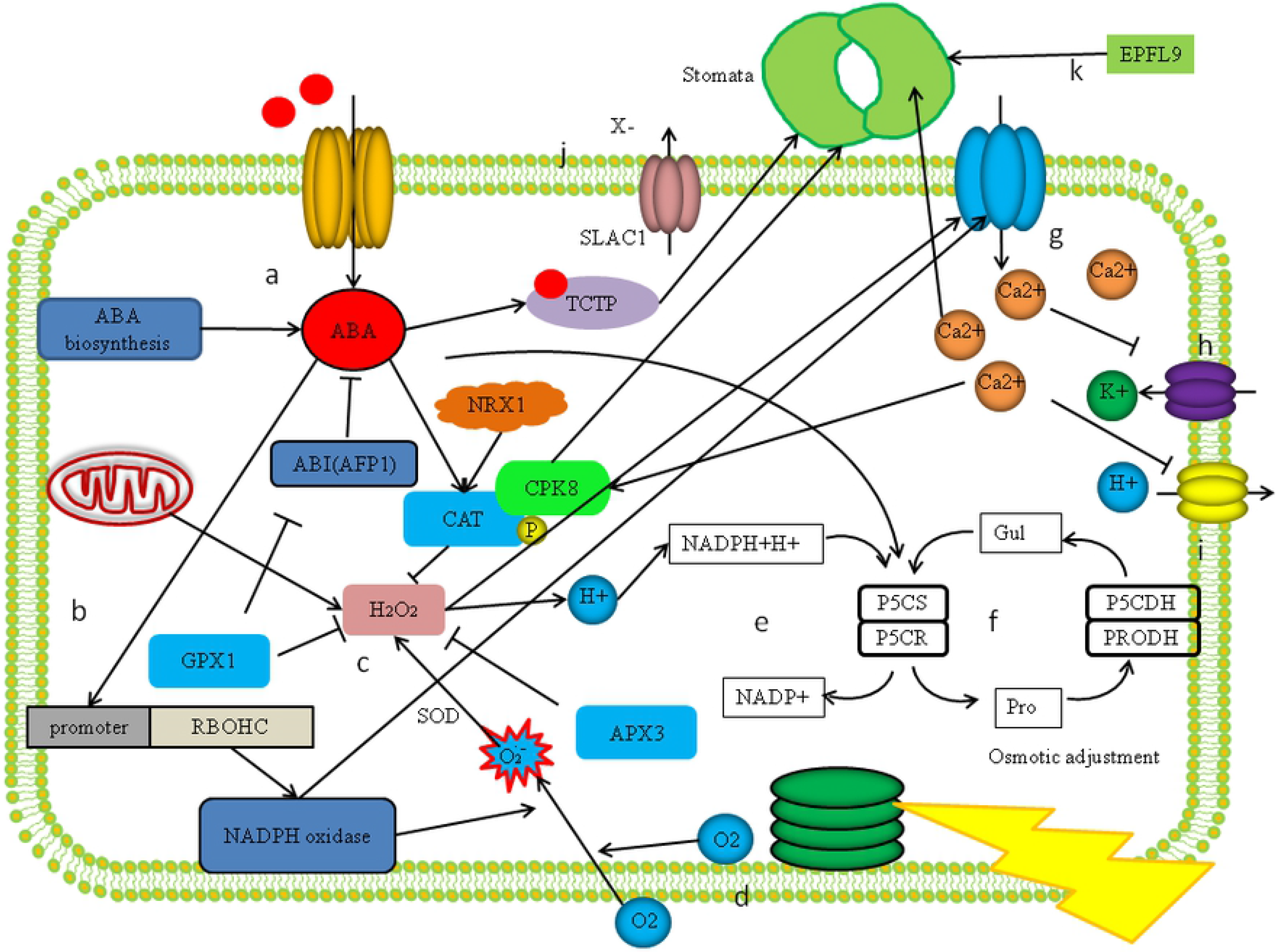
Relationship between plant antioxidant system and stomatal aperture under drought conditions.

Under drought, mitochondrial respiration can produce ROS. Oxygen produced by chloroplast photosynthesis and external oxygen are also sources of ROS in plants (Figure 6). Reactive oxygen species can destroy cell membranes and interfere with normal growth of plants, while the accumulation of ROS is toxic to plants. ABA binds to the promoter of RBOHC and promotes the production of respiratory burst oxidase (NADPH oxidase) [26]. NADPH oxidase (NOX) also activates Ca^2+^ channels on the cell membrane [27]. In addition, it has been shown to play an important protective role in plant drought stress, preventing leaves from being destroyed by ROS [28, 29]. The production of ROS activates the plant’s antioxidant system. SOD, POD, CAT, APX, GPX and other enzymes can convert superoxide anion to H_2_O_2_ and eventually decompose it into non-toxic H_2_O and O_2_ [28, 30]. The antioxidant system decomposes ROS to produce H+, which provides a substrate for glutamate synthesis in the proline pathway. At the same time, plants can also synthesize ABA under drought stress, and ABA can promote the synthesis of proline from glutamate, which eventually leads to a large accumulation of proline [31]. Proline is an important osmotic adjustment substance in plants, so the above reaction is beneficial, allowing plants to cope better with drought.

Under drought stress, the ABA signaling pathway and antioxidant system of plants are activated, and there is close interaction between them [32, 33]. On the one hand, ABA can promote the synthesis of CAT and improve the efficiency of the antioxidant system. While, on the other hand, GPX1 can inhibit ABI, thereby relieving the inhibition of ABA by plants [34]. The accumulation of ROS can activate the Ca^2+^ channel on the cell membrane, causing a large amount of Ca^2+^ to enter the cell, and so can increase the Ca^2+^ concentration of the guard cells and change their osmotic potential [35]. At the same time, high concentration of Ca^2+^ can suppress the input of K^+^ and the output of H^+^. In addition, SLAC1 transports anions out [36], resulting in an increase in the concentration of cations in the membrane. The combination of ABA and TCTP can induce stomatal closure, and CPK can phosphorylate CAT as well as promote stomatal closure [37].

EPFL9 is a gene related to stomatal development. Our results show that its relative expression decreased under drought stress, which indicates stomatal development slows or stops under drought stress. During the scavenging of H_2_O_2_ by antioxidant enzymes, it is susceptible to oxidative stress, resulting in reduced clearance. NRX1 is able to reduce oxidized antioxidant enzymes and has a stable antioxidant system [38]. However, in the gene expression level study, the expression level of *NRX1* was down-regulated. It is concluded that drought interfered with the reduction of NRX1 against oxidase.

In summary, the plant’s antioxidant system and stomatal aperture play important roles in responding to drought stress, which helps with survival. By analyzing the regulatory relationships, we show there is a close relationship between the antioxidant system and stomatal aperture. On the one hand, the antioxidant system can act as a signal to regulate stomatal aperture, while, on the other, stomatal closure can conserve water and so help maintain the stability of the antioxidant system.

## Conclusions

Drought stress can induce changes in the levels of many antioxidants in *Z. bungeanum*. Levels of POD, CAT, APX, proline and MDA all increased gradually under continuing drought, while that of SOD decreased. The expression levels of genes related to antioxidants was consistent with the changes in antioxidant content. Drought also activated genes relating to stomatal aperture, the ABA signaling pathway, the MAPK signaling pathway and the JA signaling pathway.

Under continuing drought stress the stomata of *Z. bungean*um closed gradually. The relationship between stress duration and stomatal aperture was best fitted by the exponential equation: Y=9.1529e^−0.081X^, R^2^=0.92.

## Acknowledgements

The authors would like to thank Yao Ma for his participation in the manuscript discussion. This study was financially supported by the National Key Research and Development Program Project Funding (2018YFD1000605).

